# Vaccine-induced antibodies are sufficient to limit *Salmonella* infection in the absence of complement or macrophages

**DOI:** 10.1101/2025.08.27.672591

**Authors:** Marisol Perez-Toledo, Kubra Aksu-Istil, Edith Marcial-Juarez, Ruby R Persaud, Areej Alshayea, Sian E. Jossi, Agostina Carestia, Fien von Meijenfeld, Marina Botto, Leo C James, Baas Sureward, William H Horsnell, Constantino Lopez-Macias, Ian R Henderson, OptiVaNTS consortium, Craig N Jenne, Adam F Cunningham

**Affiliations:** Institute of Immunology and Immunotherapy, University of Birmingham, Birmingham, United Kingdom; Department of Clinical Laboratory Sciences, College of Applied Medical Sciences, King Saud University, Riyadh, Saudi Arabia; Department of Microbiology, Immunology and Infectious Diseases, Cumming School of Medicine, University of Calgary, Calgary, Canada; University Medical Center Groningen, Groningen, Netherlands; Department of Immunology and Inflammation, Centre for Inflammatory Disease, Imperial College London, United Kingdom; MRC Laboratory of Molecular Biology, Francis Crick Avenue, Cambridge, United Kingdom; Division of Immunology, Institute of Infectious Disease and Molecular Medicine, University of Cape Town, Cape Town, South Africa; Medical Research Council Centre for Medical Mycology, University of Exeter, Exeter, United Kingdom; Medical Research Unit in Immunochemistry, Specialty Hospital, National Medical Center Siglo XXI, Mexican Institute of Social Security, Mexico City, Mexico; Institute for Molecular Bioscience, University of Queensland, Brisbane, Australia

**Keywords:** *Salmonella*, antibodies, complement, intravital imaging, macrophages, neutrophils

## Abstract

Antibodies to *Salmonella* Typhimurium (STm) can protect against infection. Understanding better how antibodies, complement, and leukocytes interplay can support vaccine development. We used an Outer Membrane Vesicle (OMV) vaccine against STm to study the in vivo function of anti-STm antibodies. Using intravital microscopy, we found that upon challenge, OMV-specific antibodies promote STm uptake by spleen and liver macrophages, with neutrophils rarely capturing the bacteria. Clodronate-liposome depletion of monocytic cells reveals that these cells help prevent antigen dissemination. After vaccination and challenge of C1q, C3, C4, and C5-deficient mice, all mice except C3-deficient were protected by OMV immunization. C3-deficient mice failed to mount significant germinal center and plasma cell responses; however, after the adoptive transfer of immune sera, they had lower bacterial burdens than controls. In vitro, we showed that antibodies enhance bacterial capture in macrophages. Thus, antibodies alone are sufficient to reduce bacterial burdens, but they cooperate with complement and macrophages to maximize their functions in vivo.

## Introduction

Infectious diseases continue to be a leading threat to human health. In 2019, approximately 13.7 million deaths were caused by 33 bacterial pathogens, making bacterial infections the second leading cause of death worldwide. Infections caused by *Salmonella*, including *Salmonella* Typhi and non-typhoidal *Salmonella* (NTS), were among the top 14 causes of infection-related deaths worldwide.^1^ NTS infections typically lead to self-limiting diarrhea; however, invasive (i) NTS infections are more severe and are often associated with high mortality rates in at-risk groups, such as infants in sub-Saharan Africa and individuals with HIV.^2^ Currently, there are no licensed vaccines against iNTS, although vaccine candidates are being assessed in clinical trials, including those based on outer membrane vesicles (OMV). ^3,4^ Understanding how immune responses to these types of vaccines work may support efforts to develop vaccines for *Salmonella* and other invasive Gram-negative bacteria.

Antibodies play a crucial role in the protection provided by many vaccines, and in certain situations, they also act as correlates of protection. For *Salmonella*, targeting surface antigens by antibodies, such as the lipopolysaccharide O-antigen or porins, is sufficient to reduce bacterial infection in animal models.^5–7^ Furthermore, antibodies against STm O-antigen strongly correlate with protection against iNTS infections in humans.^8^ Mechanistically, antibodies, phagocytic cells, and complement can aid in bacterial killing in vitro.^9^ However, in vitro, opsonophagocytic uptake of *Salmonella* by leukocytes can occur more rapidly than complement-mediated, cell-independent killing.^10^ This suggests that when both cell-dependent and cell-independent mechanisms of bacterial control are present, the cell-dependent pathway may serve as the primary mechanism through which antibodies exert their function. Nevertheless, antibodies and complement-derived opsonins can engage Fcγ receptors and complement receptors on immune cells, thereby enhancing phagocytosis.^11^ Thus, it is probable that both complement and cells collaborate with antibodies to optimize bacterial control. It remains to be determined whether the cooperative function of anti-*Salmonella* antibodies with phagocytic cells and complement also occurs in vivo.

In this work, we explored the requirements of selective factors associated with vaccination-induced, antibody-mediated protection in vivo. To do this, we immunized and challenged mice with an OMV-based vaccine from *Salmonella* Typhimurium (STm) and subsequently challenged them with STm. In this model, antibodies are essential for mediating protection. ^12^ By employing a combination of intravital and other imaging approaches, we find that the critical factor in the protection provided by immunization is the presence of antibodies and that multiple pathways are active, enabling these functions primarily to control bacterial numbers and localization throughout the host.

## Results

### STm preferentially associates with macrophages in the spleen and liver, regardless of vaccination

Mice immunized with outer membrane vesicles (OMV) exhibit lower bacterial burdens in the spleen and liver after challenge with STm, with antibodies playing a key role in this (Figure S1).^12,13^ To evaluate which leukocyte population is involved in STm uptake following vaccination and challenge, spleen and liver tissue sections were stained for STm, macrophages (F4/80^+^), and neutrophils (Ly6G^+^). In the spleen, both non-immunized and immunized mice showed that STm was restricted to the red pulp, with bacteria mostly associated with F4/80^+^ cells (Fig. 1A and quantification in 1B). Similarly, in the liver, STm was found to be primarily associated with F4/80^+^ cells (Fig. 1C and quantification in 1D). Therefore, STm associates with macrophages in the spleen and liver, regardless of immunization status.

**Figure 1.**
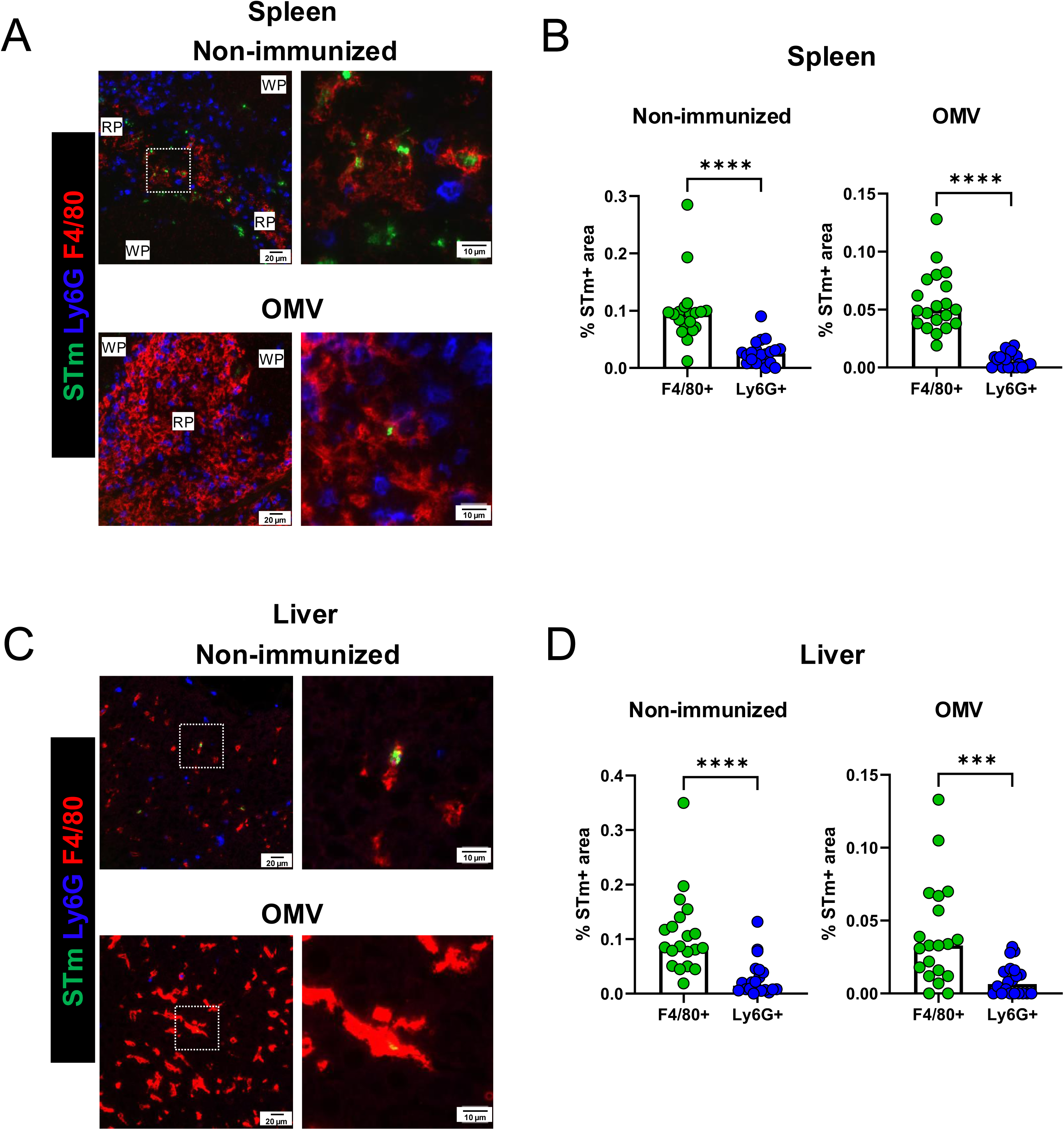
STm preferentially associates with macrophages in the spleen and liver, regardless of vaccination. (**A**) WT were immunized i.p. with 1 μg of OMVs (n=4) and challenged for 24 hours with STm SL3261 i.p. Non-immunized controls were infected alongside (n=4). Spleen and liver microsections were stained for F4/80 (red), Ly6G (blue), and STm (green). Arrows indicate positive bacteria staining. The graphs on the left show the percentage of STm positive pixel area associated with F4/80^+^ or Ly6G^+^ cells. Each dot represents a different field of view, taken from 4 different sections (OMV n=4, NI n=4). Mann-Whitney U-test. ****P<0.0001, ***P<0.0005.

### Vaccination promotes bacterial capture in the first minutes of infection

Our results at 24 hours post-challenge suggested that F4/80^+^ cells were the primary cell type involved in STm uptake after infection, with minimal involvement of neutrophils. However, this could be the result of more efficient bacterial clearance by neutrophils, which would not be detected in our analysis after 24 hours of challenge. To understand the earliest interactions between STm, macrophages, and neutrophils, intravital microscopy experiments were conducted whereby immunized and non-immunized mice were challenged with using GFP-expressing STm. In both non-immunized and immunized mice, bacteria were detected throughout the spleen and liver 10 minutes post-infection and could be observed as early as 30 seconds after challenge (Fig. 2A and 2B and videos S1-S4). In immunized mice, the frequency of immobilized bacteria over time was higher compared to non-immunized controls (Fig. 2C). Moreover, in immunized mice, bacteria showed a lower median speed, suggesting that more bacteria were captured (Fig. 2E). Additional assessments revealed that most bacteria associated with F4/80^+^ cells rather than Ly6G^+^ cells after infection (Fig. 2F), but this association with F4/80^+^ cells was most significant in vaccinated mice (Fig. 2F). The association of STm with F4/80^+^ cells showed a positive correlation with the anti-OMV IgG titers (Fig. 2G). The association of bacteria with F4/80^+^ cells was also observed at an intermediate 6-hour time point (Fig. S2 and videos S5-S6). Thus, OMV vaccination promotes bacterial uptake by F4/80+ cells, which correlates with anti-STm antibody titers.

**Figure 2.**
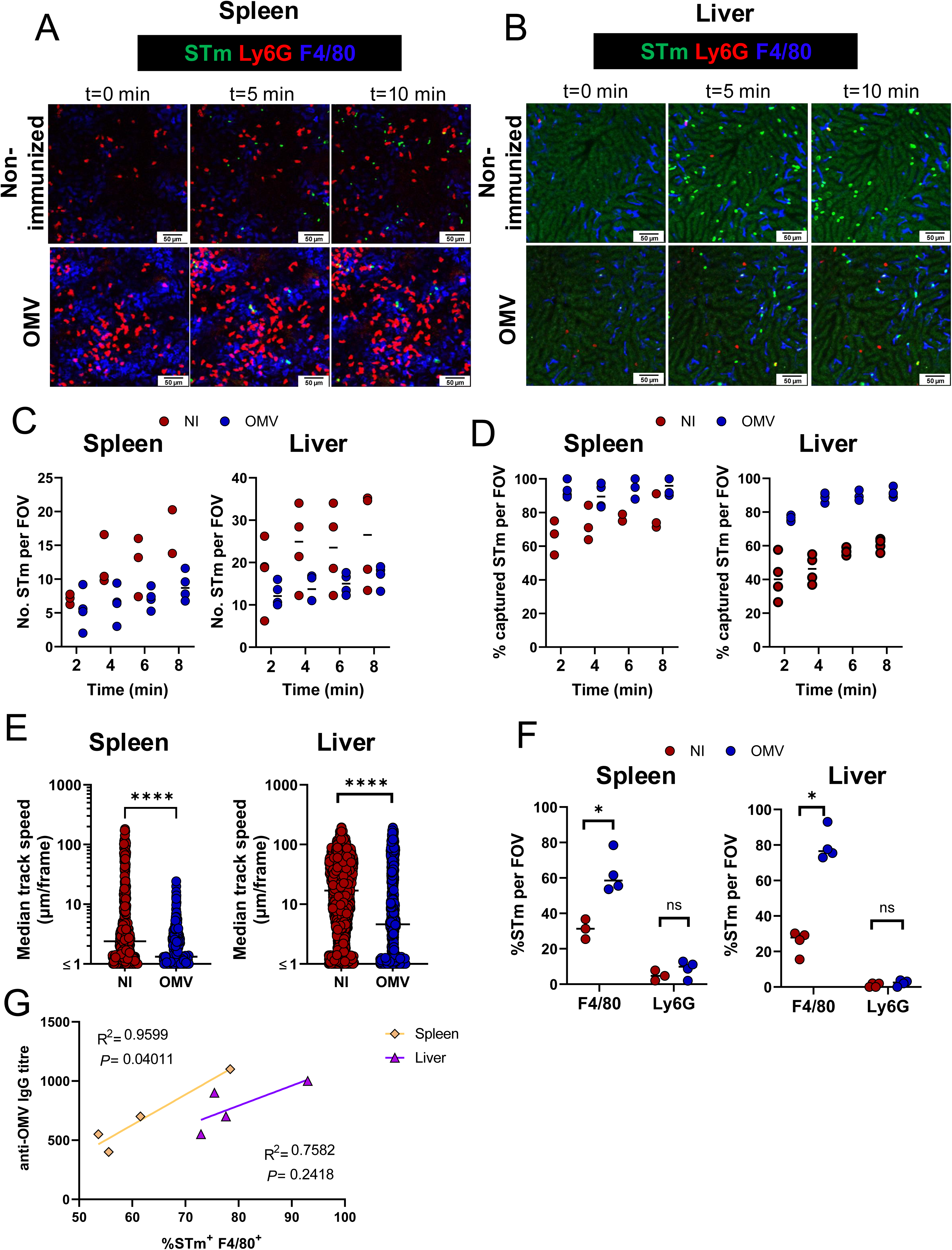
Vaccination enhances the capture of STm by F4/80^+^ cells in the spleen and liver. WT OMV-immunized and non-immunized controls (NI) were surgically prepped for live imaging to expose the liver or the spleen. 30 minutes prior imaging, anti-F4/80 and anti-Ly6G were injected i.v. When recording started, 10^7^ CFU of GFP-expressing STm were injected i.v., and videos were recorded for 10 minutes. Representative images of spleen (**A**) and liver (**B**) at the start of the recording (t=0), 5, and 10 minutes after injection of GFP-STm. Green=STm, Red=Ly6G, blue=F4/80 (**C**) Number of STm per field of view at the indicated time points post-challenge. (**D**) Frequency of captured STm per field of view at the indicated time points post-challenge. (**E**) Median track speed of STm in spleens and livers of non-immunized (NI) and OMV immunized mice (OMV). Each point represents an individual track (**F**) Frequency of STm associated with splenic F4/80^+^ cells and Ly6G^+^ cells 10 min after STm injection. (**G**) Spearman’s correlation of anti-OMV IgG titres and frequency of STm associated with F4/80^+^ cells in the spleen and liver. Mann-Whitney U-test. ****P<0.0001,*P<0.05

### Immunization-derived antibodies are sufficient to reduce bacterial burdens in the absence of monocytic cells

After establishing that macrophages are the primary cell type in the spleen and liver that capture STm, we examined the consequences of bacterial capture following the depletion of this cell population. To do this, mice were immunized with OMVs, and 24 hours prior to the challenge, macrophages were depleted using clodronate liposomes. Surprisingly, OMV-vaccinated mice treated with clodronate liposomes exhibited similar bacterial burdens to those treated with PBS liposomes (Fig. 3A). Moreover, clodronate treatment did not alter the anti-STm IgG levels (Fig. 3B). The distribution of bacteria in the spleen was then examined using immunohistology. In both non-immunized and OMV-immunized mice treated with control liposomes, bacteria were found associated with macrophages in the red pulp and were rarely observed in the white pulp (Fig. 3C). In clodronate liposome-treated, OMV-vaccinated mice, STm was abundant in the white pulp, typically associated with the follicular dendritic cell network (Fig. 3C). In the liver, bacteria were associated with macrophages in untreated and OMV-immunized mice treated with control liposomes. After clodronate treatment and the consequent absence of monocytic cells in the liver, STm remained detectable in this organ (Fig. 3C). Thus, STm capture by splenic and liver macrophages is not necessary for bacterial control but helps restrict dissemination after vaccination.

**Figure 3.**
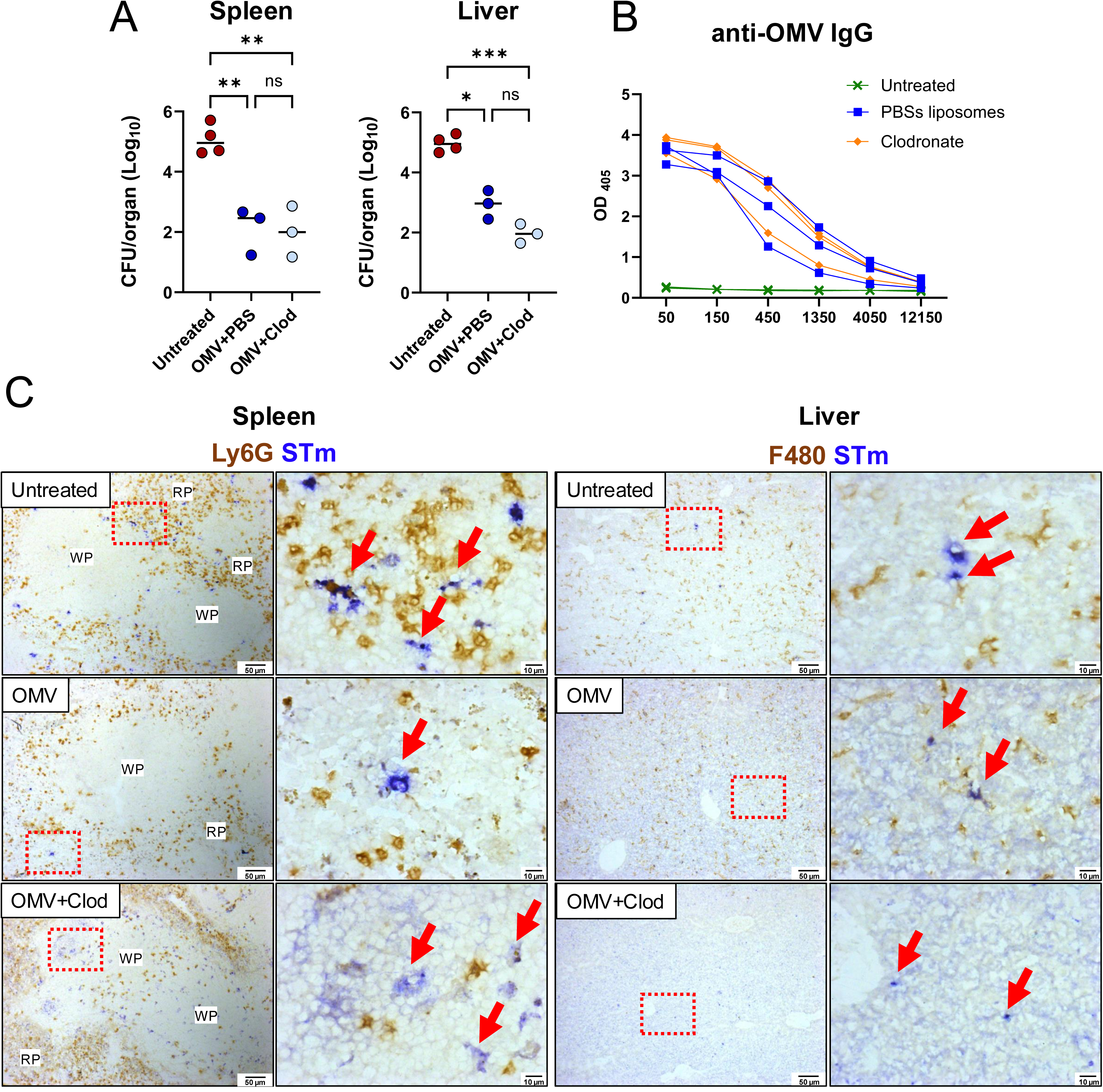
Immunization-derived antibodies are sufficient to reduce bacterial burdens in the absence of monocytic cells. WT mice were immunized with 1 μg of STm OMVs. 12 days later, mice were treated with clodronate or PBS liposomes and challenged with STm for 24 hours. (**A**) Spleen and liver bacterial burdens in STm-infected only (Untreated, n=4), OMV-vaccinated and STm-infected (OMV, n=3), and OMV-vaccinated, clodronate-treated, and STm-infected (OMV+Clod, n=3). (**B**) Anti-OMV IgG in sera from mice in (**A**) (**C**) Spleen and liver sections were stained by immunohistochemistry to detect STm (blue), Ly6G (brown) or F4/80 (brown). Arrows indicate positive STm staining. The images on the right are magnifications of the area inside the red-dotted box. WP=white pulp, RP=red pulp. One-way ANOVA, ****P<0.0001, ***P<0.0005, **P<0.005, *P<0.05

### C1q, C4 and C5, but not C3, are redundant for antibody induction to OMV

In vitro, mouse complement does not kill *Salmonella.*^14^ However, C3 has been identified as playing a role in infection control in mice in vivo.^7^ To examine these potentially conflicting roles of complement in greater depth, mice deficient in specific complement components (C1q, C3, C4, or C5) were immunized and challenged with STm for 24 hours. OMV-immunized C1q, C4, and C5-deficient mice exhibited reduced bacterial burdens in the spleen and liver compared to their non-immunized controls (Fig. 4A, B). In contrast, the bacterial burden in the spleens and livers of C3-deficient mice was similar between the immunized and non-immunized groups, indicating that the protection afforded by immunization with OMV was lost in the absence of C3 (Fig. 4A, B). To assess how these different complement-deficient mice responded to OMV and whether the reduced bacterial control in C3 mice was associated with failure to induce antibodies or a loss of functionality in antibody responses, the anti-LPS and anti-OmpD antibodies were measured. C1q, C4, and C5 all produced significant IgM and IgG responses to these antigens, whereas in C3-deficient mice, these responses were absent or minimal (Fig. 4C-F).

**Figure 4.**
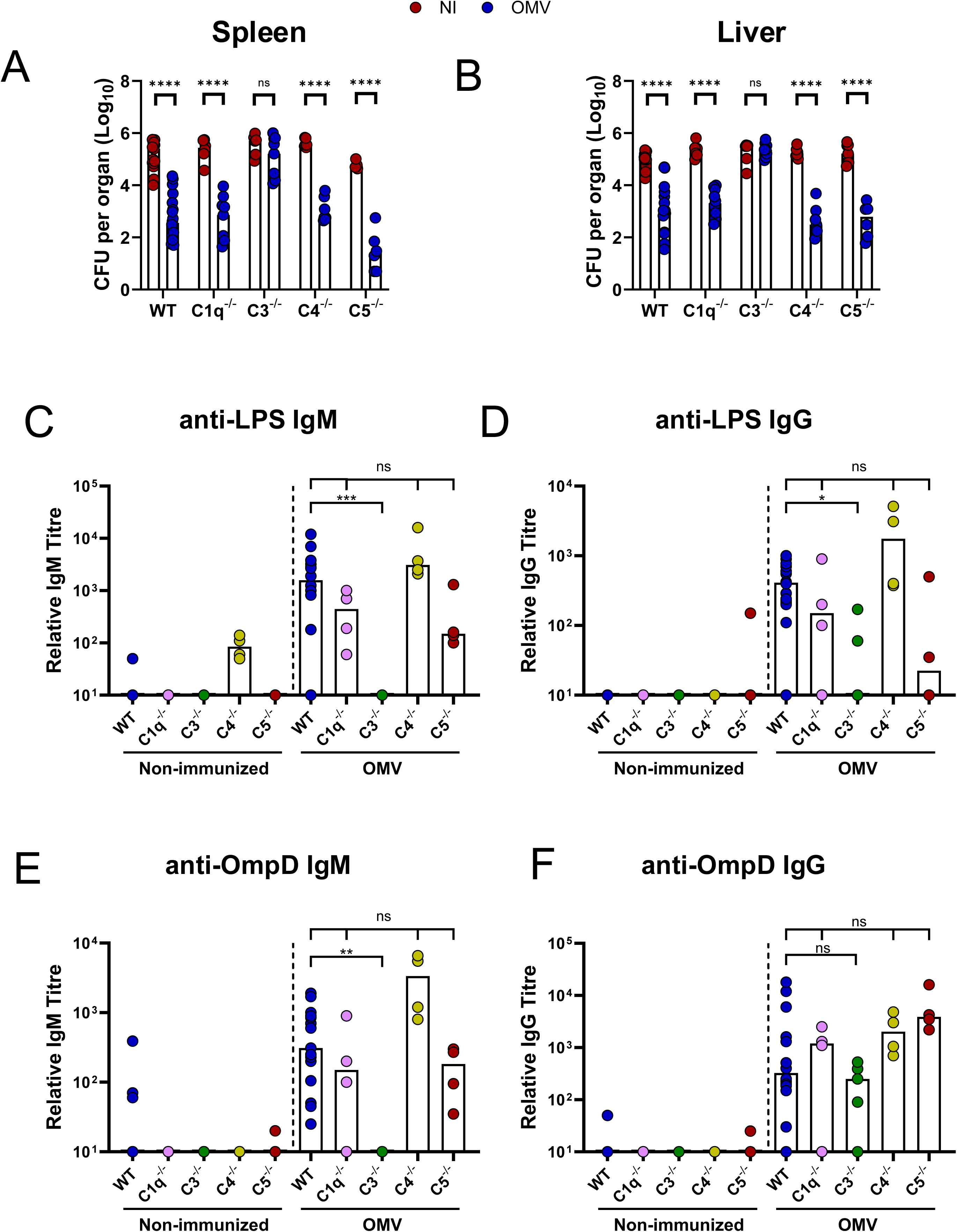
C1q, C4 and C5, but not C3, are redundant for antibody induction to OMV. C1q^-/-^(n=6), C3^-/-^(n=6), C4^-/-^(n=6), C5^-/-^ (n=5), and WT (n=11) mice were immunized with 1 µg of OMVs i.p. On day 14, they were all challenged with STm for 24 hours. Bacterial burdens in (**A**) spleen and (**B**) liver. Serum anti-STm LPS IgM (**C**) and IgG (**D**), serum anti-STm OmpD IgM (**E**) and IgG (**F**). Each point represents an individual mouse. The bars depict the median. Non-immunized C1q^-/-^n=6, C3^-/-^ n=6, C4^-/-^ n=6, C5^-/-^ n=5, and WT n=11. OMV-immunized C1q^-/-^ n=9, C3^-/-^ n=9, C4^-/-^ n=7, C5^-/-^ n=6, and WT n=18. Mann-Whitney U-test. *P<0.05, **P<0.005, ***P<0.0005, ****P>0.0001, ns= non-significant.

### Anti-STm antibodies reconstitute bacterial control in the absence of C3

The impaired antibody responses in C3-deficient mice suggested a defect in B cell responses to OMV immunization. To test this, B cell responses in the spleens of WT and C3-deficient mice were analyzed. After OMV immunization, no significant increase in the frequency or total numbers of germinal center B cells (GC) and plasma cells in C3-deficient mice was observed (Fig. 5A-D). These results suggest that C3 is necessary for the development of protective antibody responses to OMV. To determine if OMV-specific antibodies could reconstitute bacterial control in C3-deficient mice, we infected WT and C3-deficient mice with STm opsonized with either non-immune or anti-OMV-immune sera (Fig. 5E-F). WT mice infected with bacteria opsonized with OMV-specific sera had approximately a median 30-fold lower bacterial burden in the spleen and a 100-fold lower bacterial burden in the liver compared to mice infected with STm opsonized with control sera. In C3-deficient mice, the equivalent median differences in the spleen were approximately 3-fold, and in the liver, it was 4-fold (Fig. 5F). Opsonization experiments inherently involve lower antibody concentrations, as the sera are diluted 1:100 before use. Thus, we questioned whether C3-deficient mice could control the infection better if antibody levels were less limited. To investigate this, we adoptively transferred non-diluted, heat-inactivated sera from OMV-immunized mice into C3-deficient mice and infected them 24 hours later. In this case, the median fold reduction was 100-fold in the spleen and 25-fold in the liver. Taken together, our results show that C3 can contribute to the antibody-mediated control of STm infection in vivo but plays a diminishing role when antibody levels are less limited.

**Figure 5.**
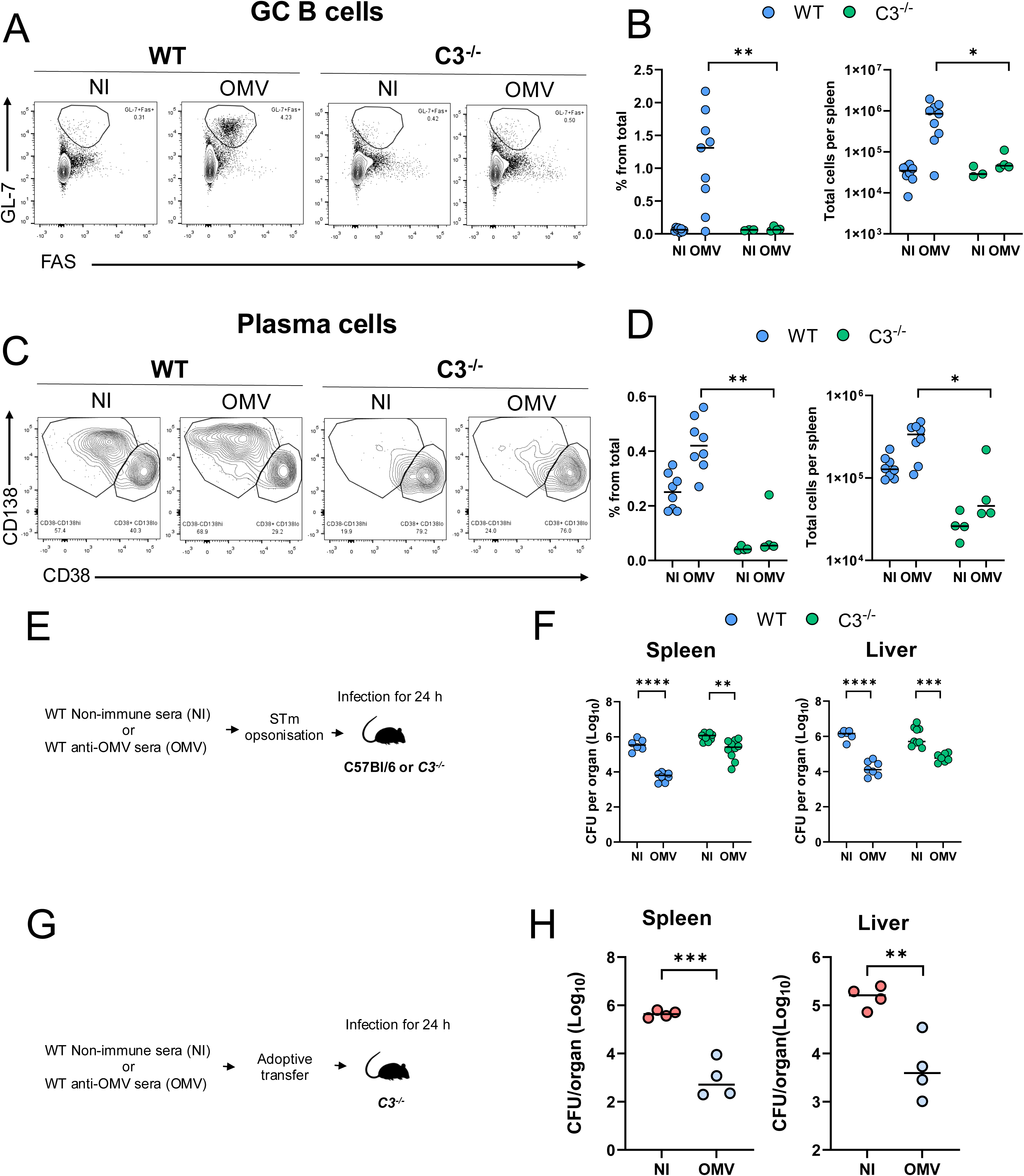
Anti-STm antibodies reconstitute bacterial control in the absence of C3. C3^-/-^ mice and WT controls were immunized with 1 μg of STm OMVs i.p. 14 days later, splenocytes were processed for flow cytometry. (**A**) Representative FACS plots of germinal centre B cells (GL-7^+^FAS^+^) in non-immunised (NI) or OMV-immunised mice. WT NI n=8, C3^-/-^ NI n=3, WT OMV n=9, C3^-/-^ OMV n=4 (**B**) Frequency and total numbers of GC B cells per spleen from (**A**). (**C**) Representative FACS plots of plasma cells (CD138^+^CD38^-^). (**D**) Frequency and total numbers per spleen of plasma cells from (**C**).WT NI n=8, C3^-/-^ NI n=3, WT OMV n=9, C3^-/-^ OMV n=4 . WT and C3^-/-^ mice were infected i.p. for 24 hours with 5 × 10^5^ CFU opsonized with non-immunized or anti-OMV mouse serum. (**E**) Experimental diagram for panel F. (**F**) Bacterial burdens in spleen and livers from (**E**). NI→WT n=6, OMV→WT n=8, NI→C3^-/-^ n=9, OMV→C3^-/-^ n=10. C3^-/-^ mice were injected intravenously with 100 μL of non-immunized (NI) or anti-OMV sera. Mice were infected i.p. with STm for 24 hours. (**G**) Experimental diagram for panel H (**H**) Bacterial burdens in spleen and liver. NI n=4, OMV n=4. Each point represents an individual mouse. The horizontal line depicts the median. B and D: Two-tailed Mann-Whitney U-test. F and H: Two-tailed unpaired t-test. ****P<0.0001 ***P<0.0005, **P<0.001

### Anti-STm antibodies and complement cooperate to enhance bacterial uptake in a human cell line in vitro

To examine whether the features associated with murine macrophages in vivo can also be observed in human-derived cells in vitro, we infected differentiated THP-1 cells with STm in the presence or absence of human serum. This takes advantage of the observation that nearly all human adults possess bactericidal antibodies against *Salmonella* ^15,16^. All sera used in these experiments had bactericidal activity that was lost after absorbing STm-specific antibodies after mixing with whole STm (Fig. 6A-B). Infection of THP-1 cells with bacteria opsonized with complete sera (CS) containing both complement and antibodies led to the highest uptake of STm (Fig. 6C). When sera that were heat inactivated (HIS) to destroy complement but had antibodies were used then bacterial uptake was increased, but this was not as high as for CS. In contrast, no increase in bacterial numbers was detected when AS was used, indicating that complement alone is not sufficient to promote bacterial uptake in this system (Fig. 6C). To test which specific complement component was required, we performed the same experiment using human serum deficient in C1, C3, or C5. Anti-STm IgG antibody levels in these sera were comparable to those observed in sera obtained from healthy volunteers (Fig. 6D). CS and C5-deficient sera promoted phagocytosis to similar levels. In contrast, opsonophagocytosis with C1q-or C3-deficient sera was lower than that with CS (Fig. 6E). This finding is consistent with the mouse studies, which suggest that antibody is sufficient. Still, that complement increases the efficiency of bacterial uptake.

**Figure 6.**
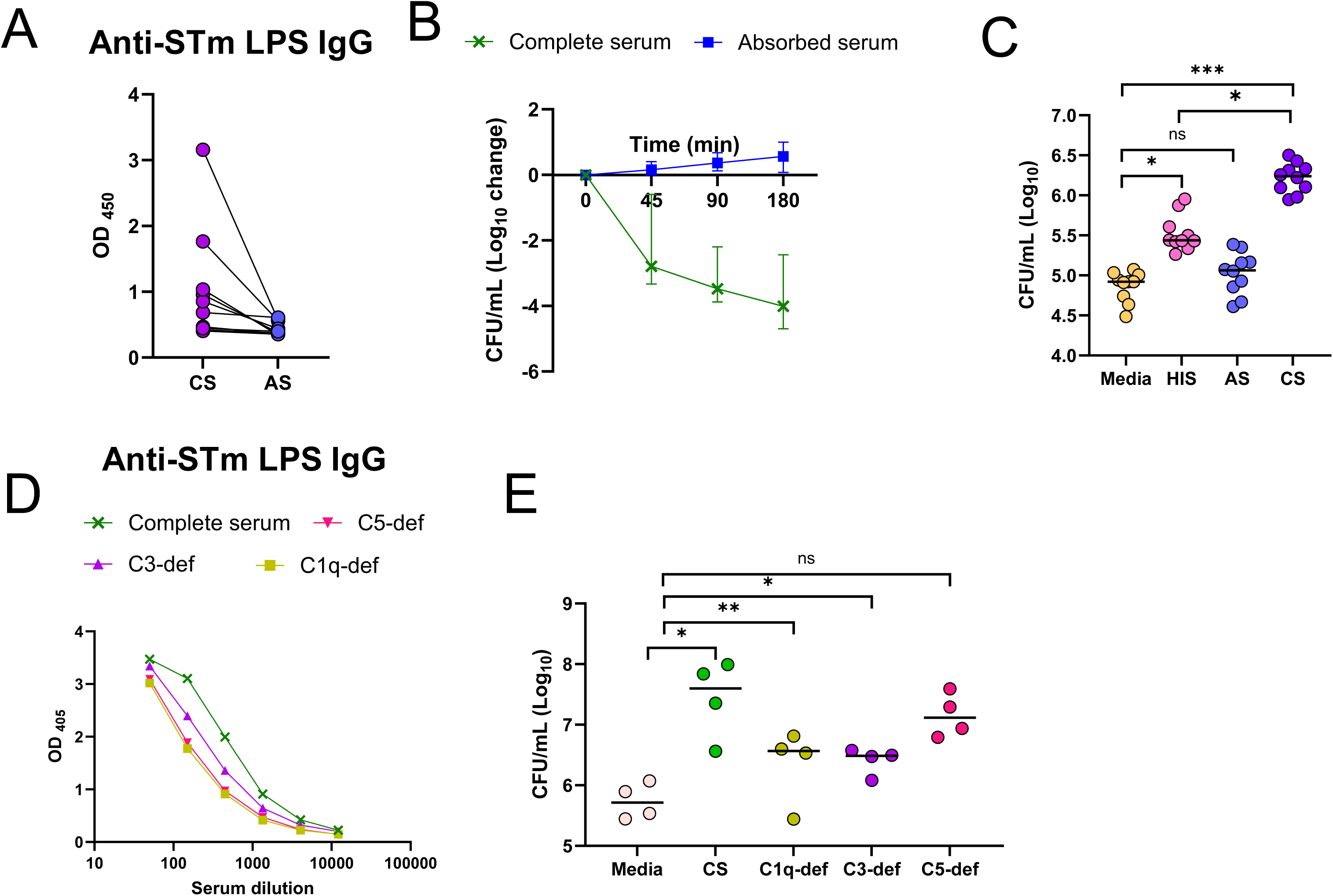
The complement system cooperates with antibodies to promote bacterial uptake by THP-1 cells. (**A**) Anti-STm LPS IgG in sera from healthy individuals in normal serum (complete serum, n=10) and after absorption with STm (absorbed serum, n=10). Sera were diluted 1:120. (**B**) Serum Bactericidal assay of complete serum (n=10) or absorbed serum (n=10) from (A). (**C**) Intracellular STm in PMA-derived THP-1 cells after incubation for 1 hour with STm and media only, heat-inactivated sera (HIS), absorbed sera (AS) or complete sera (CS), n=10. Each point represents a different donor. Horizontal lines depict the median. Further experimental details are illustrated in figure S3 (**D**) Anti-STm LPS IgG in commercial sera deficient from C1q, C3 or C5. (**E**) Intracellular STm in PMA-derived THP-1 cells after incubation for 1 hour with media only, complete sera (CS), C1q-deficient sera, C3-deficient sera or C5-deficient sera. Each point represents and independent experiment, n=4. Two-tailed one-way ANOVA. ***P<0.0005, **P>0.05, *P>0.05, ns=non-significant.

Therefore, in vitro, antibodies are sufficient to enhance bacterial uptake by THP-1 cells, while complement also plays a role in promoting this activity.

## Discussion

Antibodies play a crucial role as effector molecules for OMV vaccines.^13,17,18^ Dissecting the in vivo mechanisms through which antibodies function is made complicated by the intricate dynamics between pathogens and various components of the immune system. Our study aimed to explore the interactions among the pathogen, antibodies, phagocytes, and the complement system both in vivo and in vitro to identify the key mechanisms that lead to pathogen control. Our findings demonstrate that monocytic cells phagocytose *Salmonella* and that antibodies and complement work together to enhance this process. However, the most critical point from these studies is that antibodies can be sufficient for protection, while complement and other factors support the downstream functions that result from pathogen binding by antibodies. The intrinsic redundancy between these mechanisms underscores the importance of antibody binding for protection and provides avenues to counteract the effectiveness of pathogen escape strategies. This ultimately benefits the host by increasing the likelihood that antibodies will reduce the pathogen burden and prevent disease from developing.

Antibody activity against bacterial infections is often assessed by serum bactericidal activity or opsonophagocytic activity. In recent years, these approaches have been complemented by the development and use of systems serology approaches, which have provided additional and more nuanced insights into the effector functions of antibodies against bacterial pathogens. This enhanced understanding of how antibodies can act against bacteria may help capture the breadth of their activity against specific pathogens. This diversity in antibody function is reflected in the current study in the assessment of responses across multiple organs, both in the presence and absence of selective cell types and complement components. In vivo, a striking finding was that macrophages remained the most efficient cell type at capturing STm in the presence of antibodies. This capture by macrophages was observed in both the spleen and liver, with vaccination enhancing the efficiency of bacterial phagocytosis. Moreover, our infection studies using antibodies and THP-1 cells also demonstrated that the antibodies enhanced the efficiency of bacterial uptake into THP-1 cells in vitro. This suggests a functional overlap in these activities between mice and humans in macrophage-lineage cells.

After challenging vaccinated and clodronate-liposome-treated mice, there was a more widespread staining pattern for bacteria. Antigen was readily detected in the splenic white pulp and was particularly associated with the FDC network. This differs markedly from previous reports, where the antigen is mainly detected in the red pulp areas.^19^ Despite this, clodronate-treated mice had similar overall bacterial burdens as control mice. This suggests that a significant contribution of macrophages to the control of infection is to contain and limit bacterial spread. This is a vital role of antibodies since controlling bacterial dissemination is a key factor in limiting the risk of severe complications from infections, such as sepsis. For instance, after primary *Salmonella* infection, cardiovascular complications such as thrombosis in the spleen and liver occur in infected mice; however, we have not observed this complication in vaccinated and challenged mice. Moreover, by restricting bacterial dissemination, macrophages could help reduce the opportunities for bacteria to reach niches where they can persist and drive inflammation.

Antibody-dependent neutrophil phagocytosis activity is associated with anti-*S*. Typhi Vi responses after vaccination^20^, which led to our original hypothesis that in tissues, neutrophils are the primary cells that phagocytose STm after immunization. However, this was not the case. Intravital studies indicated that the predominant localization of bacteria to macrophages was not due to neutrophil uptake immediately after infection. Therefore, neutrophil cell death concurrent with phagocytosis of STm does not appear to contribute to the predominance of STm in macrophages following challenge of vaccinated mice. The reasons for why macrophages are the predominant host cell type associated with bacterial uptake post-vaccination in this model remain unclear. Nonetheless, this predominance does not stem from the inability of neutrophils to be infected by *Salmonella*. Previous studies, including those from our own group, have demonstrated that purified neutrophils can be efficiently infected in vitro ^21^. Moreover, we and others have demonstrated that the depletion of neutrophils has a minimal impact on bacterial control after systemic infection with STm ^19,22^. Therefore, the role of neutrophils post-vaccination in this model remains unclear. We hypothesize that macrophages outcompete neutrophils in capturing STm, leaving a limited number of bacteria in extracellular spaces available for neutrophils to phagocytose.

Meanwhile, vaccine-induced antibodies enhance the efficiency of phagocytosis by macrophages, further reducing the opportunities for neutrophils to participate in phagocytizing STm. This hypothesis will be tested in future studies.

The C1q, C4, and C5 components of the complement system were not essential for antibody-mediated protection. Non-immunized C1q-deficient mice exhibit a transient increase in bacterial numbers during the initial days of primary STm infections, but no further role for C1q in the antibody-mediated control of *Salmonella* infection has been previously assessed.^23^ Since C1q and C4 are essential components in the classical pathway for generating the C3 convertase, this suggests that the classical pathway is not crucial for reducing bacterial numbers in this model. Additionally, complement effector functions mediated by C5 are not essential for controlling *Salmonella* infection in vaccinated mice, as C5-deficient mice produced normal levels of antibodies and managed to control the infection. Since immune mouse serum does not induce cell-independent bactericidal activity against *Salmonella*, it suggests that antibodies must be functioning through alternative mechanisms to exert bacterial control ^14,24^.

Immunized C3-deficient mice were not protected against infection, which was related to their poor antibody responses to OMV, as they failed to elicit normal serum antibody, plasma cell, and germinal center responses. Nevertheless, after the adoptive transfer of immune serum into C3-deficient mice and subsequent challenge, protection was observed. Therefore, while C3 is necessary for B cell responses to develop against protective antigens within OMVs, it is not essential for the in vivo functionality of anti-STm antibodies in mice. Previously, the capture of antibody-opsonized *E. coli* by Kupffer cells in the absence of C3 has been reported, but the mechanism through which this occurs remains elusive ^25^. Work by Rossi and colleagues demonstrated a role for C3 in antibody-mediated protection after the adoptive transfer of a monoclonal IgG2a antibody to the STm LPS O4 antigen. IgG2a is the mouse IgG isotype that is most efficient at fixing complement and for FcγR-mediated mechanisms of killing. In contrast, antibody responses to OMV consist of specific antibodies of multiple isotypes against multiple antigens, which may compensate for the need for C3 in providing protection. Alternatively, C3 may become more important at different stages post-vaccination, such as in the longer term when antibody levels are lower than those observed in systems such as those used in the current study. Indeed, multiple isotypes can contribute to antibody-mediated control of STm infections in mice, as demonstrated by studies with the adoptive transfer of monoclonals and studies with IgG1-deficient mice ^6,26,27^. Thus, when multiple antigens are targeted, there may be greater redundancy for complement.

Ultimately, our study reveals that multiple mechanisms exist through which antibodies protect against *Salmonella* infections in vivo. It is likely that antibody responses to antigenically complex pathogens, such as *Salmonella*, operate through numerous, redundant, and overlapping mechanisms simultaneously. This suggests that, mechanistically, protection is associated with a signature instead of a single pathway, which could have implications for developing correlates of protection for vaccines against bacterial infections.

## Consortia

The members of the OptiVaNTS consortium (Optimising Vaccination for iNTS disease in Africa) are Galit Alter, Vassily Bavro, Joshua AJ Burton, Johanna Dean, Melita A Gordon, Ryan P. McNamara, Agnes E. Lakudzala, Mehak Zahoor Khan, Laura Fontana, and Tonney Nyirenda. Full affiliations are listed in Supplemental Table II.

## Acknowledgements

The authors would like to thank the staff from the Biomedical Services Unit of the University of Birmingham. We acknowledge the support of the University of Birmingham - Microscopy Facility, RRID: SCR_027108, and the University of Birmingham - Flow Cytometry Facility, RRID: SCR_027107, for providing access to equipment and technical expertise. We also want to thank Dr. Marina Vaysburd for their support in coordinating with reagents. This work was funded by The Wellcome Trust (Grant number 221431/Z/20/Z). MPT received a Global Mobility Award from the Wellcome Trust. AA received a scholarship from the Deanship of Scientific Research at King Saud University.

## Author Contributions

MPT conceptualized and performed experiments, curated and analyzed the data, wrote, and edited the manuscript. KAI, EMJ, RRP, AA, SEJ, AC, and FVM performed experiments and analyzed data. MB, LCJ, BS, WHH, and CLM provided resources. IRH reviewed and edited the manuscript. The OptiVaNTs consortium acquired funding and reviewed and edited the manuscript. CNJ provided resources, supervised the project, and reviewed and edited the manuscript. AFC conceptualized experiments, acquired funding, supervised the project, wrote, reviewed, and edited the manuscript. All authors read and approved the final draft of the manuscript.

## Declaration of interests

Authors declare no conflict of interest

## Declaration of generative AI and AI-assisted technologies in the writing process

During the preparation of this work the author(s) used Grammarly to correct grammar and improve the readability of the text. After using this tool, the author(s) reviewed and edited the content as needed and take(s) full responsibility for the content of the published article.

## Supplemental information

**Video S1.** Representative video of the spleen from a non-infected mouse. Mice were prepared for intravital microscopy as outlined in the methods section. Approximately 10^7^ CFUs of GFP-STm were injected intravenously at time zero. The time-lapse video shows the elapsed time in minutes, seconds, and milliseconds. Red= Ly6G, blue= F4/80, green= STm.

**Video S2.** Representative video of the spleen from a mouse immunized with STm OMVs. Mice were vaccinated with 1 µg of OMVs for 14 days and prepared for intravital microscopy as outlined in the methods section. Approximately 10^7^ CFUs of GFP-STm were injected intravenously at time zero. The time-lapse video shows the elapsed time in minutes, seconds, and milliseconds. Red= Ly6G, blue= F4/80, green= STm.

**Video S3.** Representative video of the liver from a non-immunized mouse. Mice were prepared for intravital microscopy as outlined in the methods section. Approximately 10^7^ CFUs of GFP-STm were injected intravenously at time zero. The time-lapse video shows the elapsed time in minutes, seconds, and milliseconds. Red= Ly6G, blue= F4/80, green= STm.

**Video S4.** Representative video of the liver from a mouse immunized with STm OMVs. Mice were vaccinated with 1 µg of OMVs for 14 days and prepared for intravital microscopy as outlined in the methods section. Approximately 10^7^ CFUs of GFP-STm were injected intravenously at time zero. The time-lapse video shows the elapsed time in minutes, seconds, and milliseconds. Red= Ly6G, blue= F4/80, green= STm.

**Video S5.** Representative video of the liver from a non-immunized mouse infected i.p. for 6 hours with GFP-STm. The time-lapse video shows the elapsed time in minutes, seconds, and milliseconds. Red= Ly6G, blue= F4/80, green= STm.

**Video S6.** Representative video of the liver from a mouse immunized with OMVs. Mice were immunized with OMVs and 14 days later challenged with GFP-STm i.p. for 6 hours. The time-lapse video shows the elapsed time in minutes, seconds, and milliseconds. Red= Ly6G, blue= F4/80, green= STm.

## Methods

### Mice, bacterial strains, immunogens and immunization

Outer Membrane Vesicles (OMV) were prepared from a TolR-deficient *Salmonella* Typhimurium strain 14028 as described previously ^13^. Protein content was quantified with a bicinchoninic acid assay kit following the manufacturer’s instructions (Thermo Scientific, Cat. No. 23227). *Salmonella* Typhimurium LPS was purchased from Enzo (Cat. No. ALX-581-011-L002). OmpD from *Salmonella* Typhimurium was purified as described previously ^5^. Wild-type (WT) C57BL/6J mice were obtained from Charles River UK (strain #27). C1q-deficient mice ^28^, C3-deficient mice ^29^, and C5-deficient ^30^ mice were kindly donated by Marina Botto. Leo C. James kindly donated C4-deficient mice. ^29^ For all experiments, a mix of male and female mice, aged 6 to 12 weeks, was used. Procedures were carried out at the Biomedical Services Unit of the University of Birmingham or the University of Calgary, and were performed with complete local and national ethical approval (UK Project License P06779746, University of Calgary Animal Protocol AC22-0057). Mice were housed in specific-pathogen-free conditions, under a 12-hour light/dark cycle, with a controlled temperature range of 20-24 °C. All mice were euthanized by cardiac bleed under anesthesia, followed by cervical dislocation.

Mice were immunized intraperitoneally with 1 µg of OMV or PBS as non-immunized controls. 14 days post-immunization, mice were injected intraperitoneally (i.p.) with 5×10^5^ CFU of a *Salmonella* Typhimurium Δ*aroA* SL3261 strain ^31^. To prepare the bacteria for challenge, one colony from STm SL3261, grown in Luria-Bertani (LB) agar, was inoculated into 10 mL of LB broth and incubated overnight at 37 °C. The following day, the bacteria were regrown in fresh LB medium, with shaking at 180-200 rpm and 37 °C, until the optical density at 600 nm was approximately 1.0. 1 mL of the culture was then centrifuged at 6000 *g* and washed twice with sterile PBS. The final pellet was resuspended in 1 mL of PBS and further diluted in PBS to achieve the final concentration for injection. Serial dilutions of the leftover bacterial preparation confirmed the administered bacterial dose. Mice were sacrificed 24 hours post-challenge. Livers and spleens were collected to quantify the bacterial burden in organs.

Approximately 1 mg of tissue was homogenized through a 70-μm cell strainer and reconstituted in 1 mL of sterile PBS. Neat and serial 1:10 dilutions were plated in LB agar plates and incubated at 37 °C overnight for CFU counting.

Clodronate-depletion experiments were performed as described previously ^32,33^. Briefly, WT mice were immunized with OMVs as described above. 12 days post-immunization, mice were injected i.v. with 200 µL of clodronate liposomes (Clodrosome®, CLD-8909) or PBS liposomes (Encapsome ®, CLD-8901, both from Encapsula NanoSciences). 24 hours later, mice were infected with STm as described above. Unvaccinated mice infected with STm that were not injected with liposomes served as untreated controls. For opsonization experiments, 5×10^5^ bacteria/mL were mixed for 30 minutes with pooled heat-inactivated anti-OMV sera (diluted 1:100), or pooled heat-inactivated non-immune sera (diluted 1:100). Mice were then infected with 5×10^5^ opsonized bacteria via i.p. Opsonized bacteria were plated onto LB agar plates to confirm that the opsonization did not affect viability. For passive transfer experiments, C3-deficient mice were injected intravenously with 100 µL of pooled heat-inactivated non-immune or anti-OMV sera, prepared from mice that had been immunized twice, 35 days apart, with 1 µg of OMV. Twenty-four hours later, the mice were infected as described above.

### Intravital Microscopy

Male C57Bl/6 mice were vaccinated with OMVs as previously described. 14 days post-immunization, the mice were anesthetized using a combination of ketamine hydrochloride (Anesketin, Dechra Veterinary Products Inc.) at a dosage of 200 mg/kg and xylazine (Nerfasin 100, Dechra Veterinary Products Inc.) at 10 mg/kg. Liver and spleen were then prepped and imaged following established protocols ^19^. Anti-mouse F4/80 (1.6 µg, clone BM8, BioLegend) and anti-Ly6G (1.6 µg, clone 1A8, BioLegend) were injected i.v. 30 min prior to imaging. Once the recording was started, mice were challenged intravenously with 10^7^ CFU of *Salmonella* Typhimurium constitutively expressing EGFP in a pDiGc plasmid ^34^. 10-minute-long videos were recorded using an inverted Leica SP8 microscope (Leica Microsystems). Image analysis was performed with Fiji (version 2.9.0).

### ELISA

96-well Nunc Maxisorb plates (ThermoFisher Scientific, Cat No.442404) were coated with either STm LPS, STm OmpD, or STm OMVs at 5 μg/mL diluted in carbonate buffer. After blocking for 1 hour at room temperature with 2% bovine serum albumin (BSA) (Sigma, Cat. No. A7906-100G), mouse serum was added starting at a 1:50 or a:100 dilution, and then further diluted three-fold. After incubation for one hour at 37 °C and washing with PBS-0.001% Tween 20, AP-conjugated anti-mouse IgM (diluted 1:2000) or IgG (diluted 1:1000) (Both from SouthernBiotech, details in Table S1) were added to the wells, followed by a further one-hour incubation at 37 °C. After washing with PBS-0.001% Tween 20, development was performed with Sigma FAST™ p-Nitrophenyl Phosphate tablets (Cat. No. N2770), prepared according to the manufacturer’s instructions, and the absorbance was measured at 405 nm approximately 1 hour later. Titers were calculated by identifying the dilution at which the OD_405_ was equal to 2.

### Sample preparation for FACS

Sample preparation for FACS analysis was performed as described before ^35^. Briefly, single-cell suspensions from spleens were obtained by mashing approximately 20 mg of tissue through 50 µm cell strainers (CellTrics Cat. No. 04-0042-2317) in 5 mL of RPMI 1640 (Gibco Cat. No. 31870-025) supplemented with 5% FBS (Gibco Cat. No. A5256801) and 5 mM EDTA (PanReac AppliChem Cat. No. A4892). Red blood cells were lysed with ACK lysis buffer (Life Technologies, Cat. No. A10492-01), and after washing, the cell suspension was adjusted to 10^7^ cells/mL. 2.5×10^6^ cells were then incubated with an anti-CD16/32 antibody (Invitrogen Cat. No. 14-01-61-85) in FACS buffer (2% FBS, 5 mM EDTA and 0.01% sodium azide). Live/Dead staining was performed with ZombieAqua^TM^ or ZombieViolet^TM^, following the manufacturer’s instructions (Biolegend Cat. No. 423101 and 423113). Cell suspensions were incubated on ice for 25 minutes with a mix of the primary antibodies in FACS buffer. Samples were acquired using a BD LSR II Fortessa flow cytometer, and data were analyzed using Flowjo Software v10.8.1 (BD Life Sciences). Detailed information about the antibodies used can be found in the Table S1.

### Immunohistology

5 µm cryosections of spleens and livers were fixed in acetone (Acros Organics Cat. No. 268310025) for 20 min and then stored at −20 °C until analysis. For immunohistochemistry, the sections were re-hydrated in Tris-buffered Saline (pH 7.6) at room temperature and stained in the same buffer with primary antibodies for 45 minutes. HRP-conjugated or biotin-conjugated secondary antibodies, along with the Vectastain® ABC-AP alkaline phosphatase kit (Vector Laboratories Cat. No. Ak-5000) were used as secondary reagents. HRP activity was detected with SIGMA*FAST* 3-3’Diaminobenzidine tablets (Sigma-Aldrich Cat. No. D4293), while alkaline-phosphatase activity was detected using naphtol AS-MX phosphate and fast blue salt, with levamisole (All from Sigma-Aldrich).

For immunofluorescence, the sections were re-hydrated in PBS (pH 7.4) and blocked for 10 minutes with 10% fetal bovine serum (FBS) in PBS. Additional biotin-blocking steps were performed for the liver prior to staining using an avidin/biotin blocking kit (Vector Laboratories Cat. No. SP-2001) following the manufacturer’s instructions. Antibodies were incubated in the dark at room temperature for 40 minutes. Slides were then mounted in Prolong Diamond (ThermoFisher Cat. No. P36970) and allowed to cure for 24 hours at room temperature in the dark before imaging. Images were acquired with a Zeiss Axio Scan Z1 slide scanner (Zeiss, Germany). A detailed list of primary and secondary antibodies used is provided in Table S1.

### Removal of anti-Salmonella antibodies from serum and serum bactericidal assay

Serum was collected from healthy individuals by venipuncture and stored at −80°C until needed (Ethical approval: ERN_1805 December 2023). C1q (Cat. No. A509), C3 (Cat. No. A508), and C5-depleted sera (Cat. No. A501) were obtained from QuidelOrtho. To remove specific STm antibodies, 900 µL of human serum was incubated with 100 µL of a hyper-concentrated suspension of washed STm D23580 at 4 °C for one hour. After centrifugation at 6000 g for 5 minutes, the supernatant was further absorbed 2 times. The absorbed serum was aliquoted and stored at −80°C until needed. For the serum bactericidal assay, approximately 10^5^ CFU of mid-log STm D23580 were incubated with complete serum or absorbed serum. The samples were incubated in a shaking incubator at 37 °C, and aliquots were taken at 45, 90, and 180 minutes. Each aliquot was diluted 10-fold and plated in LB agar plates to quantify the CFU/mL. The change in the CFU was determined by subtracting the base 10 logarithm of the CFU at the specified time point from the base 10 logarithm of the CFU at time zero.

### In vitro gentamicin protection assay with THP-1 cells

THP-1 cells were obtained from ATCC (TIB-202). Cells were cultured in RPMI-1640 complete medium containing 10% FBS (Gibco Cat. No. A5256801), 1% L-glutamine (Gibco Cat. No. A5256801), and 1% penicillin-streptomycin (Gibco 15140-122) at 37 °C with 5% CO_2_. Differentiation of THP-1 cells was performed by incubating them with 200 ng/mL Phorbol 12-Myristate 13-Acetate (Sigma-Aldrich, Cat. No. P8139) for 2 days, following by a rest period of 3 days. 2 days before the experiment, cells were washed once in PBS, resuspended at a density of 2×10^5^ cells/mL and seeded in 96-well plates. Cells were infected with STm strain D23580^36^, at a multiplicity of infection (MOI) of 10. Before infection, the bacteria were opsonized for 10 minutes in the presence of human serum at a 1:10 dilution.

The infection was performed for 1 hour, followed by a further 1 hour incubation in the presence of gentamicin (Gibco Cat. No. 15710-064) at 100 μg/mL. The bacterial count was determined by lysing the cells with 0.2% deoxycholate, followed by serial 10-fold dilutions in PBS. Dilutions were plated in LB agar plates and incubated overnight at 37 °C.

### Image analysis

Image analysis was conducted in Fiji. To determine the frequency of STm+ pixels in Ly6G or F4/80 cells, four different fields of view were captured from each spleen and liver. For each image, a Gaussian blur (sigma=1) and background subtraction were applied. Then, each image was split into green (STm), blue (F4/80), and white (Ly6G) channels. Pixels present in both STm and F4/80, or STm and Ly6G, were identified using the Boolean operator “AND” in the Image Calculator function. The resulting image was binarized with auto threshold, and the Analyze Particles function was used to quantify the pixel area.

To quantify the total number of STm per field of view, GFP-only snapshots from specific time points were taken, and the number of particles was measured using the Analyze Particles function. To determine the total number of STm captured per field of view, GFP-positive events that remained stationary (moving less than 1 cell diameter for ≥3 minutes) were manually counted. The number was expressed as a percentage, with the total number of STm at a specific time set to 100%. To assess the association of STm with either F4/80 or Ly6G cells, a snapshot of the last frame of each video was analyzed. The image was split into green (STm), blue (F4/80), and white (Ly6G) channels. Then, an image showing the pixels present in both STm and F4/80, or STm and Ly6G, was produced using the Boolean AND operator in the Image Calculator function. This resulting image was converted to binary using auto threshold, and the Analyze Particles function was used to quantify the pixel area. The results are expressed as a percentage, where the total number of STm in the same frame is set to 100%. To analyze the particle movement, time-lapse stacks were analyzed with TrackMate ^37^.

### Statistical analysis

All experiments were conducted at least twice. Group sizes were determined using G*Power 3.1.9.2 based on the expected differences between groups to achieve a study with at least 80% power and a p<0.05 significance level. The number of mice per group was assigned depending on available litter sizes and was paired to include animals of similar age whenever possible. Colony Forming Unit counts were expressed as the base 10 logarithm to approximate a normal distribution and were analyzed using a two-tailed unpaired t-test for comparisons between two groups, or a two-tailed one-way ANOVA for comparisons involving more than two groups (Figures 3A, 4A and 4B, 5F and 5H, 6C and E). The two-tailed Mann-Whitney U test was used to evaluate the significance of data from the remaining in vivo experiments comparing two groups. P-values were calculated with GraphPad Prism version 10.1.0 and were considered significant when p≤ 0.05.

### Data availability

Additional data is available from the AFC upon reasonable request.

## Notes

### Competing Interest Statement

The authors have declared no competing interest.

