## Supplemental Figures for "Vaccine-induced antibodies are sufficient to limit *Salmonella* infection in the absence of complement or macrophages"

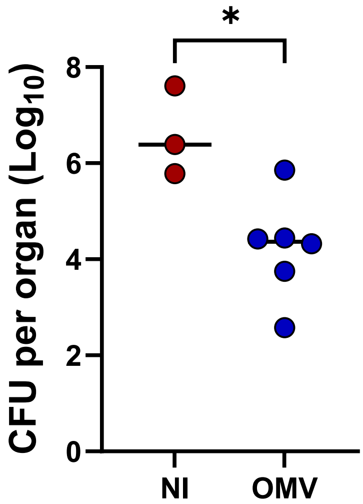


Supplemental Figure 1. Immunization with OMVs reduces the bacterial burden in the spleen upon challenge with STm LT2. WT mice were immunized with 1 µg of STm OMVs. On day 14, mice were infected i.p. with $10^7$ CFU of STm LT2 for 6 hours. Horizontal lines depict the median. Each point represents an individual mouse. Two-tailed unpaired t-test. *P<0.05.

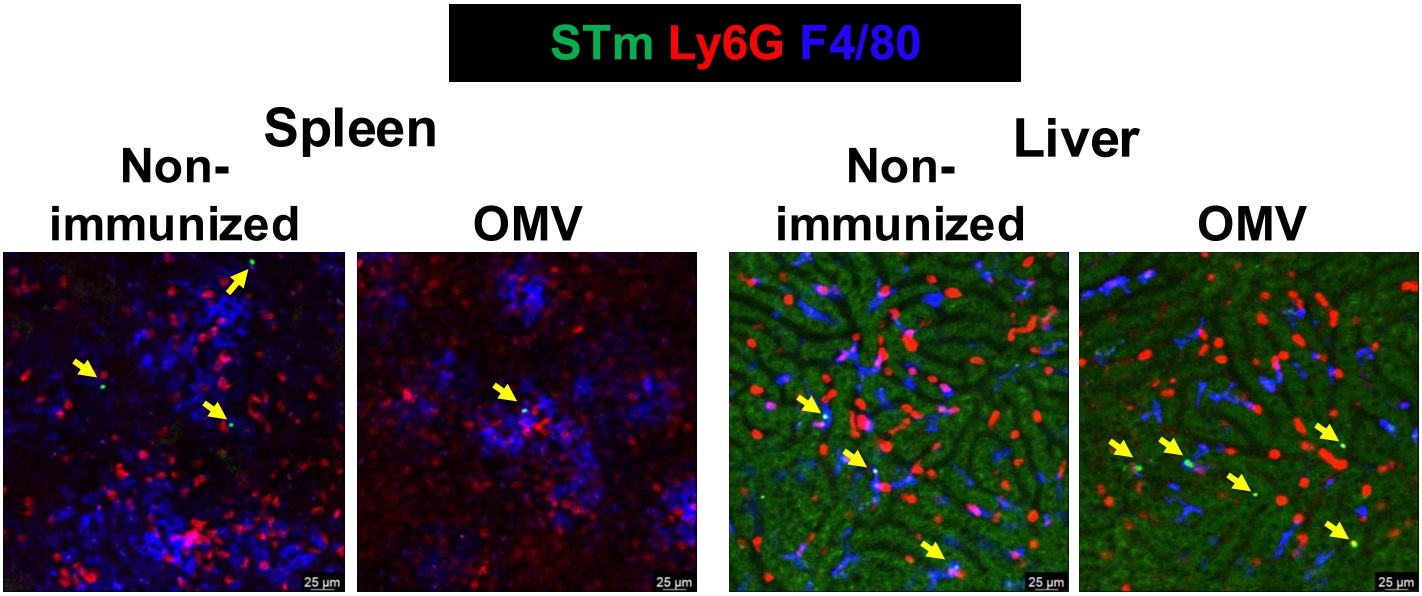


Supplemental Figure 2. STm is found associated with F4/80+ cells in the spleen and liver 6 hours post-infection, regardless of vaccination status. WT mice were immunized with 1 µg of STm OMVs. On day 14, mice were infected i.p. with $10^7$ CFU of STm LT2 for 6 hours. Mice were prepped for live imaging as described in the methods. 30 minutes prior imaging, mice were injected i.v. with anti-mouse Ly6G (red) or anti-F4/80 (blue). Yellow arrows indicate the presence of bacteria.

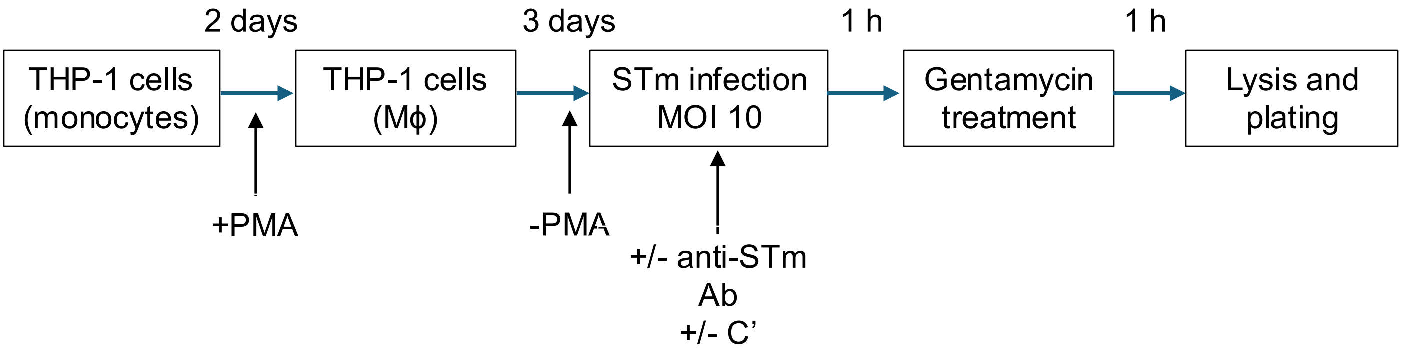


Supplemental Figure 3. THP-1 cells were differentiated with PMA for 2 days and then rested without PMA for an additional 3 days. Following this, the cells were infected with *Salmonella* Typhimurium (STm) at a specified multiplicity of infection (MOI), either in the presence or absence of anti-STm antibodies and complement. After an incubation period of 1 hour, the cells were treated with gentamicin for another hour. Subsequently, the cells were lysed, diluted, and plated for colony-forming unit (CFU) counting.
